# *In vitro* lymph node-mimicking 3D model displays long-term T cell-dependent CLL proliferation and survival

**DOI:** 10.1101/2023.04.03.535388

**Authors:** Marco Vincent Haselager, Bianca Francisca van Driel, Eduard Perelaer, Dennis de Rooij, Danial Lashgari, Remco Loos, Arnon P. Kater, Perry D. Moerland, Eric Eldering

## Abstract

Chronic lymphocytic leukemia (CLL) cells are highly dependent on microenvironmental cells and signals. The lymph node (LN) is the critical site of *in vivo* CLL proliferation and development of resistance to both chemotherapy and targeted agents. We present a new model that incorporates key aspects of the CLL LN which enables investigation of CLL cells in the context of a protective niche. We describe a 3D *in vitro* culture system utilizing ultra-low attachment (ULA) plates to create spheroids of CLL cells derived from peripheral blood (PB). Starting from CLL:T cell ratios as observed in LN samples, CLL activation was induced by either direct stimulation and/or indirectly via T cells. Compared to 2D cultures, 3D cultures promoted CLL proliferation in a T cell-dependent manner, and enabled expansion for up to 7 weeks, including the formation of follicle-like structures after several weeks of culture. Addition of LN-derived stromal cells further enhanced the proliferative capacity. This model enables high-throughput drug screening, of which we describe response to Btk inhibition, venetoclax resistance, and T cell-mediated cytotoxicity as examples. In summary, we present the first LN-mimicking *in vitro* 3D culture for primary CLL, which enables readouts such as real-time drug screens, kinetic growth assays and spatial localization. This is the first *in vitro* CLL system that allows testing of response and resistance to venetoclax and Btk inhibitors in the context of the tumor microenvironment, thereby opening up new possibilities for clinically useful applications.

## Introduction

CLL is a prime example of a malignancy that is highly dependent on interactions with the microenvironment. At LN sites, CLL cells are provided with external signals from surrounding cells such as CD40L-presenting T helper cells, myeloid and stromal cells, creating a protective niche that is thought to play a key role in the long-term response to targeted agents and development of resistance^1^. Currently, CLL biology and response to drugs are widely studied using primary leukemia cells isolated from PB, which has important drawbacks given the fact that *in vivo* CLL growth occurs at LN sites^2^. Although coculture with stromal cells or the addition of soluble factors can increase and extend survival of CLL cells, no existing systems permit the long-term expansion of CLL cells *in vitro*^*3*^. Moreover, current models cannot be used to study adhesion, migration or immune responses in the context of the tumor microenvironment. Despite similarities between human and murine models for CLL, inherent differences pose important limitations, such as the lack of LN involvement in most commonly used CLL mouse models^4^.

Development of 3D models that are able to accurately mimic the CLL microenvironment may help to bridge the gap between current *in vitro* systems and the physiologic CLL microenvironment *in vivo*, overcoming some of the current limitations of *in vitro* CLL studies. Recent efforts have been made towards the development of bone marrow-specific CLL models^5,6^, yet it is currently accepted that the LN is the critical site of CLL proliferation^2^. We attempted to establish a model that incorporates key aspects of CLL LN biology, taking into account physiological relevance. We formulated several criteria that the system should meet; contribution of non-CLL cells, long-term proliferation, induction of drug resistance, high-throughput, and practical to use.

Here, we present a novel 3D LN model for the *in vitro* culture of CLL that mimics a LN microenvironment using PB samples of CLL patients. In order to integrate multiple facets of the CLL microenvironment, we tested inclusion of non-CLL cell populations such as T cells, myeloid cells, and stromal cells. As CD4^+^CD40L^+^ T cells have been shown to cluster near Ki-67^+^ CLL cells within LN tissue biopsies^7^, we hypothesized that improved cell-cell interactions in a 3D conformation aid CLL cells to proliferate and evade cell death^3,8^. For this reason, we used patient samples containing a relatively high number of CD4^+^ T cells to mimic CD4^+^CD40L^+^ T cell help in the LN. As multiple cell types in the LN microenvironment may affect drug resistance and proliferation of CLL cells via cytokine secretion^1^, we also applied a cytokine cocktail of IL-21, IL-15 and IL-2 together with the TLR9 agonist CpG. IL-21 is produced by CD4^+^ T cells and follicular T cells, and primes CLL cells for proliferation but also renders them more sensitive to apoptosis^3^. IL-15 is produced by monocytes and may enhance CLL proliferation and survival^9^. IL-2 shares these properties and is primarily produced by CD4^+^ and CD8^+^ T cells^10,11^. Importantly, CD40 activation upregulates the IL-21, IL-15 and IL-2 receptors^12,13^, which may further prime CLL cells for proliferation and survival. Finally, we applied CpG to activate TLR signaling in CLL cells, as is shown by gene expression studies in LN-derived CLL cells^14^, and which may further promote the upregulation of cytokine receptors^15^. In this way, we created a model where multiple upstream signals activate divergent downstream signaling pathways in CLL cells, resulting in phenotypes that allow for better investigation of CLL proliferation and high-throughput drug screening.

In this paper, we demonstrate the principle of the model including several functional readouts such as proliferation, spatial localization, and drug resistance. We provide the protocols and details in order for other research groups to adopt this system. Moreover, this model can be adapted and used for the generation of more complex tumor microenvironments tailored to specific research objectives.

## Material & Methods

### Patient material

After written informed consent, patient blood samples were obtained during diagnostic or follow-up procedures at the Departments of Hematology and Pathology of the Academic Medical Center Amsterdam. This study was approved by the AMC Ethical Review Biobank Board under the number METC 2013/159 and conducted in accordance with the Declaration of Helsinki. Blood mononuclear cells of patients with CLL, obtained after Ficoll density gradient centrifugation (Pharmacia Biotech, Roosendaal, The Netherlands) were cryopreserved and stored as previously described^16^. Cells were stained with the following antibodies: CD8, CD56, CD5, CD19, TCRγδ, CD3, CD4 and CD20. Cells were measured by flow cytometry on a Canto II flow cytometer (BD Biosciences, Franklin Lakes, NJ, USA). We used CLL samples containing 50-90% CD5^+^/CD19^+^ CLL cells and >10% CD4^+^ T cells. Paired PB and LN samples were characterized in the context of the HOVON 158 clinical trial.

### Antibodies

| <u>Product</u> | <u>Article number</u> | <u>Supplier</u> |
| --- | --- | --- |
| **Patient sample characterization** |  |  |
| Anti-human CD8 | CYT-8F8 | Cytognos (Madrid, Spain) |
| Anti-human CD56 | CYT-56PE | Cytognos (Madrid, Spain) |
| Anti-human CD5 | 341109 | BD Biosciences (Franklin Lakes, NJ, USA) |
| Anti-human CD19 | IM3628 | Beckman Coulter (Indianapolis, IN, USA) |
| Anti-human TCR$\gamma\delta$ | 655410 | BD Biosciences (Franklin Lakes, NJ, USA) |
| Anti-human CD3 | 345767 | BD Biosciences (Franklin Lakes, NJ, USA) |
| Anti-human CD4 | 300524 | BioLegend (San Diego, CA, USA) |
| Anti-human CD20 | 302320 | BioLegend (San Diego, CA, USA) |
| **Flow cytometry** |  |  |
| Anti-human CD4 | 46-0047-42 | eBioscience (Santa Clara, CA, USA) |
| Anti-human CD8 | 344732 | BioLegend (San Diego, CA, USA) |
| Anti-human CD14 | 340436 | BD Biosciences (Franklin Lakes, NJ, USA) |
| Anti-human CD19 | 555412 | BD Biosciences (Franklin Lakes, NJ, USA) |
| Anti-human CD25 | R0811 | Dako (Santa Clara, CA, USA) |
| Anti-human CD56 | CYT-56PE | Cytognos (Madrid, Spain) |
| Anti-human CD95 | 305621 | BioLegend (San Diego, CA, USA) |
| **Confocal microscopy** |  |  |
| Alexa Fluor 488 anti-human CD4 | 300519 | BioLegend (San Diego, CA, USA) |
| Alexa Fluor 594 anti-human CD8a | 301056 | BioLegend (San Diego, CA, USA) |
| Alexa Fluor 647 anti-human CD14 | 325612 | BioLegend (San Diego, CA, USA) |
| Alexa Fluor 700 anti-human CD19 | 302226 | BioLegend (San Diego, CA, USA) |

#### Protocols

All used protocols are listed in *Supplemental information* together with corresponding reagents and equipment.

#### Cell culture

PBMCs of CLL patients were plated in standard round-bottom 96 well culture plates (2D) or in ULA plates and centrifuged for 10 minutes at 1000rpm and subsequently incubated for 24 hours to allow spheroid formation (3D). Cells were cultured at 37⁰C and 5% CO_2_ using standard cell culture techniques. Cells were cultured in Iscove’s Modified Dulbecco’s Medium supplemented with 10% fetal calf serum and 1% penicillin/streptomycin. CLL cells were cultured as total PBMCs or in coculture with stroma in a 5:1 ratio. Cells were stimulated as indicated: either directly with a B cell stimulation cocktail consisting of 25ng/mL IL-2, IL-15 and IL-21 including 1µg/mL CpG and/or indirectly via a T cell stimulation of anti-CD3 (clone 1XE) and anti-CD28 (clone 15E8) antibodies.

#### Primary lymph node fibroblasts

Primary lymph node fibroblasts were a kind gift of T. de Jong and L. van Baarsen (Department of Experimental Immunology, Amsterdam UMC). Inguinal lymph node needle biopsies were obtained from rheumatoid arthritis patients, healthy controls, and donors with autoantibodies at-risk for developing rheumatoid arthritis. Biopsies were enzymatically digested with dispase, collagenase, and DNAse. Afterwards, fibroblasts were grown out over the course of several weeks of *in vitro* culture.

#### Flow cytometry

For proliferation assays, PBMCs were labeled with either CFSE or CellTrace Violet. After culture, spheroids were resuspended and disintegrated to ensure proper antibody staining. Cells were incubated with monoclonal antibodies for surface staining for 30 minutes at 4⁰C. Cells were stained with the following antibodies: CD4, CD8, CD14, CD19, CD25, CD56, CD95 and a viability dye. Samples were measured on a Canto II flow cytometer (BD Biosciences, Franklin Lakes, NJ, USA). Samples were analyzed using FlowJo software.

#### Confocal microscopy

During culture, spheroids were imaged using standard light field microscopy. After culture, the supernatant was removed, and spheroids were embedded in 1% agarose to preserve spheroid structure. Agarose-embedded spheroids were then fixated in 4% paraformaldehyde for 10 minutes at room temperature, permeabilized in 0.01% Triton-X for 10 minutes at room temperature, blocked in 0.5% BSA for 1 hour at room temperature and washed with PBS in between steps. Immunofluorescent staining of spheroids was carried out via incubation with primary antibodies overnight at 4⁰C and incubation with secondary antibodies for 1 hour at room temperature while washing with PBS in between steps. Cells were stained with the following antibodies: CD4-AlexaFluor488, CD8-AlexaFluor549, CD14-AlexaFluor647 and CD19-AlexaFluor700. Afterwards, cells were stained in DAPI solution for 10 minutes at room temperature. Finally, agarose-embedded spheroids were transferred to an 18 well glass bottom microscopic slide and analyzed using an SP8-X SMD confocal microscope (Leica Microsystems, Wetzlar, Germany).

#### Kinetic growth assays

3D cultures were stimulated and treated as indicated. Culture plates were placed in an IncuCyte live-cell imager (Essen Biosciences, Royston, United Kingdom) in an incubator at 37⁰C and 5% CO_2_. Scans were taken every 6 hours using the single spheroid assay for live-cell analysis application using 4x magnification. Spheroid area was quantified using IncuCyte software as a proxy for spheroid growth.

#### Venetoclax sensitivity assay

CLL cells were cultured as indicated for 24 hours. After culture, cells were resuspended and incubated with and without venetoclax for an additional 24 hours. CLL cell viability was measured by flow cytometry using DioC6 and TOPRO-3 viability dyes. Samples were measured on a FACS Calibur flow cytometer (BD Biosciences, Franklin Lakes, NJ, USA). Samples were analyzed using FlowJo software. Specific apoptosis is defined as [% cell death in treated cells] – [% cell death in medium control] / [% viable cells medium control] x 100%.

#### T cell cytotoxicity assay

Primary CLL cells were labeled with CellTrace Violet according to the manufacturer’s instructions and cocultured with healthy donor T cells in a 4:1 effector to target ratio for 24 hours. PBMCs were incubated in the presence of either 1ng/mL blinatumomab, 10µM venetoclax, or medium. The viability of the target cells was assessed by flow cytometry using TOPRO-3 and MitoTracker Orange viability dyes. Samples were measured on an LSR Fortessa flow cytometer (BD Biosciences, Franklin Lakes, NJ, USA). Specific lysis is defined as [% target cell death in treated sample] – [% target cell death in medium control] / [100% -% target cell death in medium control] x 100%.

#### Statistics

Statistical analysis was performed using GraphPad Prism 9 software. A paired t-test was used to analyze paired observations between 2 groups. Two-way ANOVA with multiple comparisons and Tukey correction for multiple comparisons was used to analyze differences between 3 or more groups. In the case of drug titrations, Geisser-Greenhouse correction was added in combination with Tukey correction for multiple comparisons to analyze differences between groups using two-way ANOVA. *p<0.05; **p<0.01; ***p<0.001; ****p<0.0001.

## Results

### Design of a PBMC-based LN model

First, we aimed to incorporate T cells as a source of physiological CD40L by recapitulating the composition of CLL and T cells in the CLL LN. To this end, we confirmed that LN samples contained significantly more T cells compared to PB samples which was specifically due to the increased number of CD4^+^ T cells in the LN (Figure 1A-B)^17^. For this reason, we selected PB samples from our CLL biobank containing >10% CD4^+^ T cells to simulate a LN-like setting using primary PBMCs.

**Figure 1.**
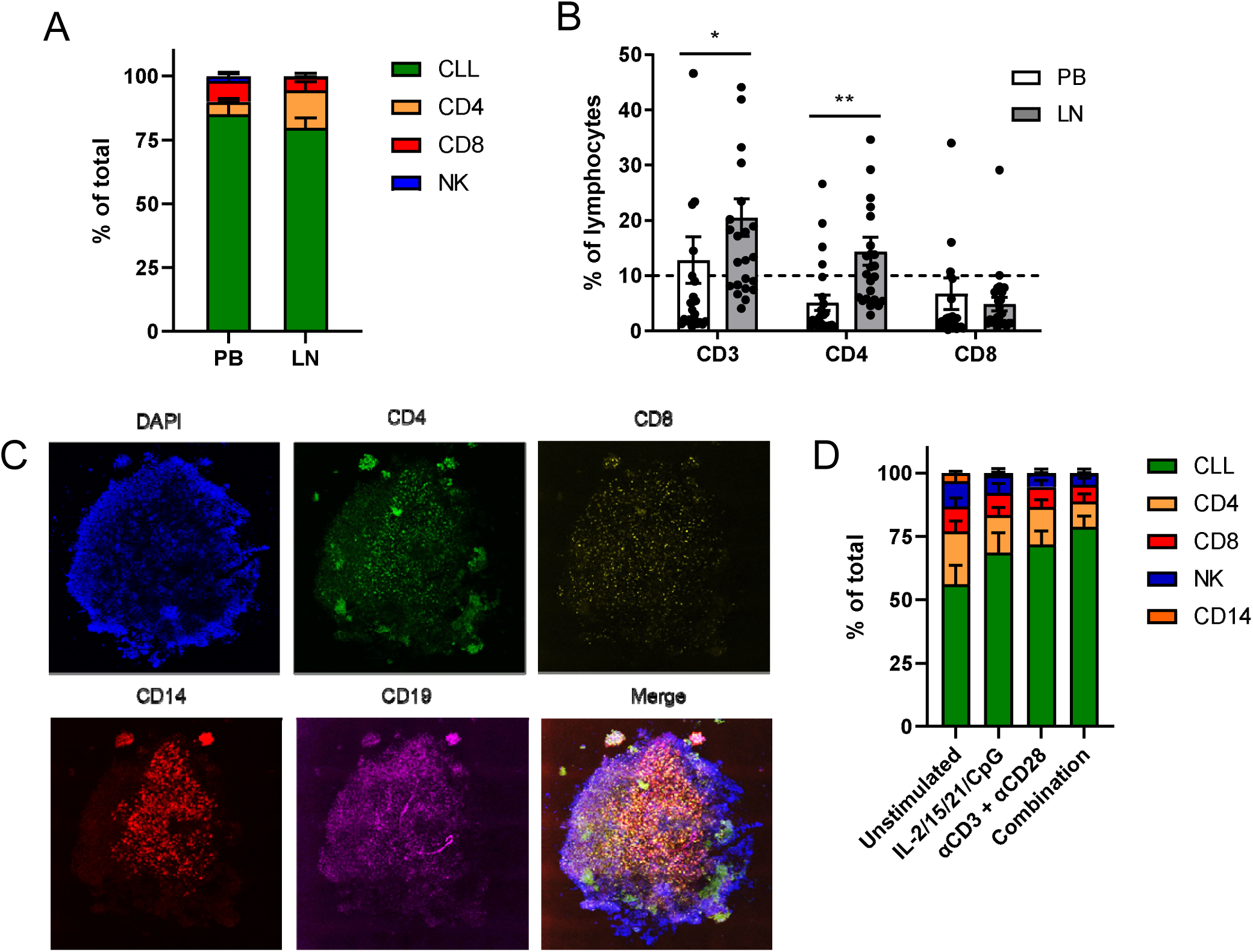
Design of a PBMC-based LN model. **A-B)** Lymphocytes were isolated from paired PB and LN samples and characterized in treatment-naïve CLL patients (n=24). **(A)** Cells were gated based on the following surface markers: CLL (CD5+/CD19+), CD4^+^ T cells (CD3+/CD4), CD8^+^ T cells (CD3+/CD4+), and NK cells (CD56+). Healthy B cells (CD5-/CD19+) were excluded from the analysis due to very small populations. Error bars represent the mean ± SEM. **(B)** Detailed analysis of the T cell subsets. The dashed line represents the >10% CD4^+^ T cell cut-off we applied in selection of PB biobank samples. Error bars represent the mean ± SEM (n = 24), *p<0.05, **p<0.01 (two-way repeated measures ANOVA). **C)** CLL sample after spheroid formation. The sample was stained for DAPI (blue), CD4 (green), CD8 (yellow), CD14 (red), and CD19 (magenta) and analyzed by confocal microscopy. **D)** CLL samples were stimulated as indicated and cultured in ULA plates for 5 days. Cells were gated based on the following surface markers: CLL (CD5+/CD19+), CD4^+^ T cells (CD3+/CD4+), CD8^+^ T cells (CD3+/CD4+) and NK cells (CD56+). Healthy B cells (CD5-/CD19+) were excluded from the analysis due to very small populations. Error bars represent the mean ± SEM (n=8).

After spheroid formation and self-organization of cells, we observed several follicles of CD4^+^ T cells in both the spheroid center and periphery, a population of CD8^+^ T cells that was evenly distributed throughout the spheroid and a network of CD14 myeloid cells around the spheroid center (Figure 1C). To further characterize the model, we investigated the cell composition of the CLL spheroids after 5 days of *in vitro* culture upon different modes of stimulation. We optimized a cocktail independent of CD40 stimulation to directly stimulate the CLL cells, consisting of IL-2, IL-15, IL-21 and CpG Supplemental Figure 1). Alternatively, we applied T cell stimulation using αCD3 and αCD28 antibodies. Whereas unstimulated spheroids contained approximately 50% CLL cells and 20% CD4^+^ T cells, upon either mode of stimulation the CLL fraction expanded to approximately 70% (Figure 1D). Consistently, the combination of both stimulations resulted in a further expansion of CLL cells. In order to keep the system as simple as possible and as complex as necessary, for most readouts presented here we only applied IL-2/15/21/CpG stimulation unless indicated otherwise.

### 3D cultures promote CLL proliferation in a T cell-dependent manner

As a marker for responsiveness of CLL cells to soluble stimuli and cell-cell interactions within 3D cultures, we measured expression of CD95. Stimulation with IL-2/15/21/CpG induced maximal CLL activation (Figure 2A). T cell stimulation via αCD3/αCD28 antibodies also resulted in maximal activation of CLL cells, thereby demonstrating the interactions of CLL and T cells in the model. Yet, T cells required stimulation with αCD3/αCD28 antibodies for 100% activation (Figure 2B). Direct stimulation of CLL cells with IL-2/15/21/CpG resulted in an induction of CLL proliferation with 60% dividing CLL cells in 2D cultures and 80% in 3D cultures (Figure 2C). Stimulation of T cells via αCD3/αCD28 antibodies also triggered CLL proliferation with 40% dividing CLL cells, but dividing cells only proliferated once or twice (Figure 2D). Combination of these two modes of stimulation resulted in approximately 70% dividing CLL cells in both 2D and 3D cultures, yet 3D cultures showed a significantly increased proliferation index upon stimulation in contrast to 2D cultures (IL-2/15/21/CpG: p = 0.0297; combination: p = 0.0012) (Figure 2E). This is a result of most cells proliferating once or twice in 2D cultures, whereas 3D cultures showed a similar number of cells in each generation (Figure 2D). Analysis of T cell proliferation shows that only direct T cell stimulation via αCD3/αCD28 antibodies resulted in cell division of all T cells in addition to a significantly increased proliferation index (Supplemental Figure 2).

**Figure 2.**
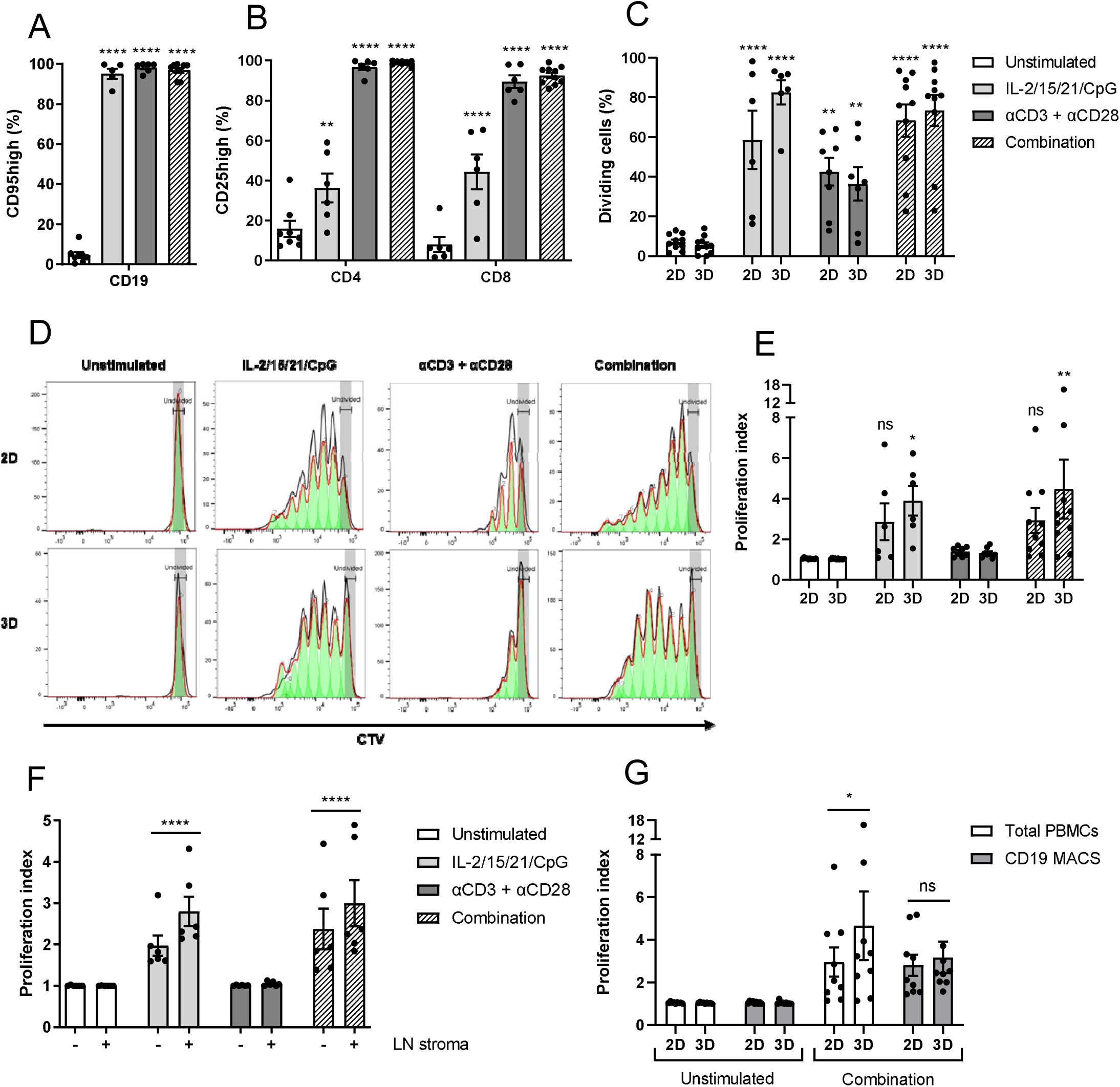
3D cultures promote CLL proliferation in a T cell-dependent manner. **A-B)** CLL samples were stimulated as indicated and cultured in ULA plates for 5 days. Afterwards, cells were measured using flow cytometry with the CD95 activation marker for CLL cells **(A)** and CD25 as the activation marker for CD4^+^ or CD8^+^ T cells **(B)**. Error bars represent the mean ± SEM (n = 10), **p<0.01, ****p<0.0001 (Figure A: paired t-test; Figure B: two-way ANOVA). Each group was compared with the unstimulated control group. **C-E)** CTV-labeled CLL samples were stimulated as indicated and cultured in standard round-bottom 96 well plates (2D) or ULA plates (3D) for 8 days and measured using flow cytometry **(C-D)**. Afterwards, the proliferation index was quantified **(E)**. Bars represent the mean ± SEM (n = 10), *p<0.05, **p<0.01, ****p<0.0001 (two-way ANOVA). Each group was compared with the unstimulated control group. **F)** CTV-labeled CLL cells were stimulated as indicated and/or cocultured with primary LN fibroblasts in 3D spheroid cultures for 1 week. Afterwards, cells were measured using flow cytometry, and the proliferation index was quantified. Error bars represent the mean ± SEM (n = 6), *p<0.05 (two-way repeated measures ANOVA). **G)** Stimulation of IL-2/15/21/CpG and αCD3/αCD28 antibodies was performed on the total pool of PBMCs and in parallel on the CD19^+^ CLL fraction without the presence of T cells. The proliferation index was quantified. Error bars represent the mean ± SEM (n = 9), *p<0.05 (two-way repeated measures ANOVA). MACS, magnetic-activated cell sorting.

To investigate the incorporation of additional cell types into the model, we performed experiments using LN-derived fibroblasts obtained from non-CLL donors. Without stimulation, LN fibroblasts did not induce CLL proliferation, but in combination with IL-2/15/21/CpG stimulation, LN fibroblasts increased proliferation in every individual CLL sample (Figure 2F). These results indicate that addition of LN stromal cells may further promote CLL proliferation, suggesting that the introduction of additional cell types may further influence the culture of CLL cells. Due to the scarcity of primary LN material, we further employed the model here without the addition of LN fibroblasts. Finally, we investigated the contribution of T cells to the proliferation of CLL cells. The significantly increased proliferation index observed in 3D cultures was abrogated in the absence of T cells (Figure 2G). These data suggest that 3D cultures promote CLL proliferation via T cells.

### 3D cultures enable the long-term proliferation of CLL cells with the emergence of follicle-like structures

Next, we investigated the longevity of CLL cultures after stimulation with IL-2/15/21/CpG. For this purpose we did not apply αCD3/αCD28 antibody stimulation, as then T cells rapidly proliferate and dominate the culture. After 1 week of culture, stimulated 3D cultures expanded in a linear fashion with a 2-fold expansion at 2 weeks of culture, whereas cell numbers in 2D cultures started to collapse and eventually reached a plateau (Figure 3A). In the 3D cultures, a second expansion phase occurred at around 4 weeks. Interestingly, spheroids started to develop follicle-like structures around 3-4 weeks of culture (Figure 3B). We observed that T cells localized both at the spheroid core as well as in these follicle-like structures (Figure 3C). Additionally, CLL cells in the spheroids were predominantly CXCR5^+^ whereas CLL cells at the core and follicles were CXCR5^-^, indicating multiple subpopulations of CLL cells. Furthermore, follicle-like structures showed an enrichment of Ki-67^+^ CLL cells, suggesting that these are also sites of proliferation (Figure 3D). These data show that 3D cultures enable the expansion and long-term proliferation of CLL cells, with emergence of self-organizing, follicle-like structures.

**Figure 3.**
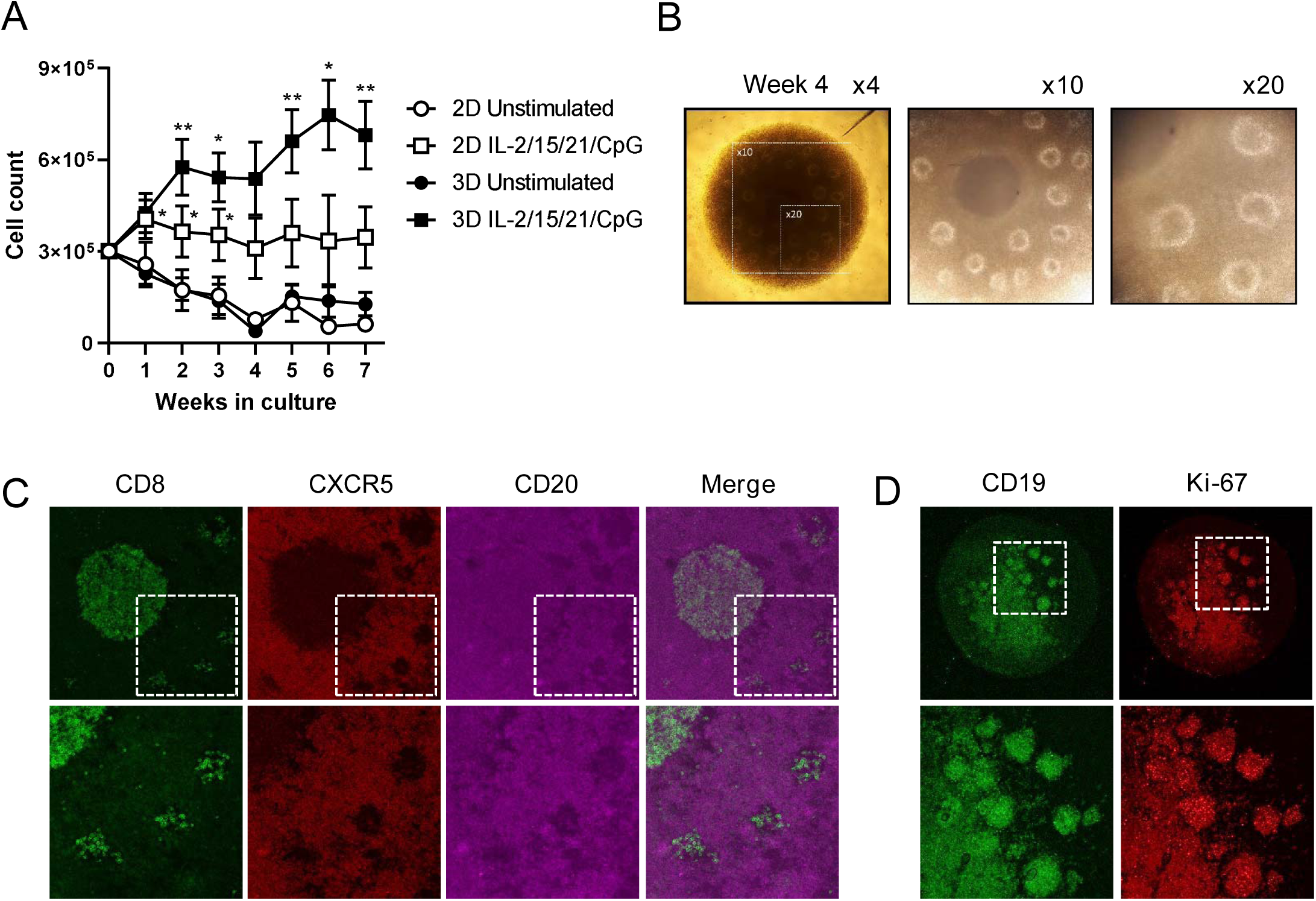
3D cultures enable the long-term proliferation of CLL cells. **A-D)** CLL samples were stimulated as indicated and cultured in standard round-bottom 96 well plates (2D) or ULA plates (3D) for 7 weeks. **A)** Total cell counts were measured every week by flow cytometry. Error bars represent the mean ± SEM (n = 6), *p<0.05, **p<0.01 (paired t-test). Each group was compared with the unstimulated control group. **B)** Spheroid morphology observed by light field microscopy. The panels show 4x, 10x, and 20x magnifications of the spheroid at week 4. **C)** Spheroids were cultured for 4 weeks and stained for CD8 (green), CXCR5 (red), and CD20 (magenta). Spheroids were imaged by confocal microscopy, including magnifications in the bottom panel. **D)** Spheroids were cultured for 4 weeks and stained for CD19 (green) and Ki-67 (red). Spheroids were imaged by confocal microscopy, including magnifications in the bottom panel.

### Btk targeting inhibits spheroid growth and impacts spheroid architecture

A 3D model enables novel assays and readouts such as kinetic growth assays using live-cell imaging. We performed a proliferation experiment in the presence of the Btk inhibitor acalabrutinib and imaged spheroids in real-time from which the spheroid area was quantified as a proxy for spheroid growth. Independent of the presence of acalabrutinib, spheroids stimulated with IL-2/15/21/CpG showed an initial growth to 4 × 10^6^ µm^2^ over the course of 3 days (Figure 4A). Though untreated spheroids reached a plateau in spheroid growth, acalabrutinib treatment decreased spheroid size in a dose-dependent manner, eventually reaching the size of unstimulated spheroids upon 0.1-1µM acalabrutinib. To test the accuracy of this assay as a readout for cell proliferation, we subjected the imaged spheroids to standard flow cytometry as an endpoint readout. Based on CellTrace Violet signal, the proliferation index was quantified in each spheroid, confirming significantly decreased proliferation (1µM acalabrutinib: p = 0.0348) in a dose-dependent manner (Figure 4B-C). Though both IncuCyte and flow cytometry readouts were consistent, the IncuCyte assay provided not only real-time data, but also seemed more sensitive compared to flow cytometry. Notably, acalabrutinib treatment affected not only spheroid size, but also impacted spheroid architecture (Figure 4D). As a proxy for cell-cell adhesion, we plotted the area of the spheroid core as a percentage of total spheroid area, where a lower percentage translates to more detached cells. Stimulation of spheroids with IL-2/15/21/CpG resulted in the formation of a dense spheroid core structure consisting of adherent cells, expanding to approximately 70% of the total spheroid area as a result of proliferation. Acalabrutinib treatment was associated with a lower core expansion (30-40%) and increased areas of low-density regions located at the outer spheroid edges, consisting of detaching cells that can easily be dissociated (Figure 4E).

**Figure 4.**
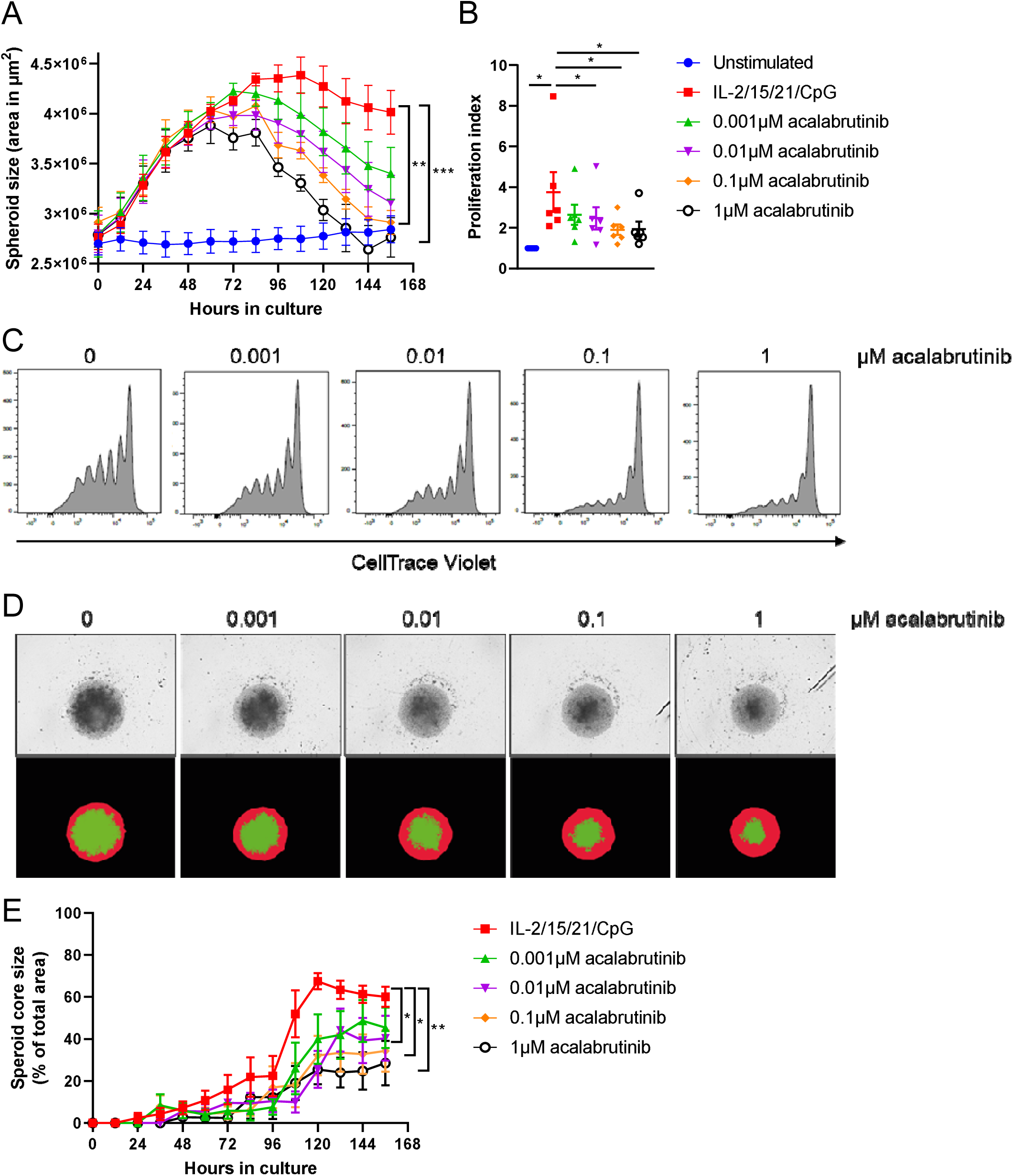
Btk targeting inhibits spheroid growth and impacts spheroid architecture. **A-C)** CTV-labeled CLL samples were stimulated with IL-2/15/21/CpG and cultured in ULA plates (3D) for 6 days in the presence of 0.001-1µM acalabrutinib. Culture plates were placed in an IncuCyte live-cell imager which imaged and quantified the spheroid area every 12 hours **(A)**. Error bars represent the mean ± SEM (n = 6), **p<0.01, ***p<0.001 (two-way ANOVA). Afterwards, cells were measured using flow cytometry **(C)**, and the proliferation index was quantified **(B)**. Error bars represent the mean ± SEM (n = 6), *p<0.05 (paired t-test). **D-E)** Based on the live-cell imaging data, spheroid morphology was analyzed by quantifying the total spheroid area (red mask) and quantifying the spheroid core area (green mask) **(D)**. The spheroid core area was then plotted over time as a percentage of the total spheroid area, where 100% equals to an identical area of red and green masks **(E)**. Error bars represent the mean ± SEM (n = 6), *p<0.05, **p<0.01 (two-way repeated measures ANOVA).

### CLL spheroids enable the study of drug resistance and T cell cytotoxicity

Furthermore, this 3D CLL model can be used to investigate both *in vitro* and *in vivo* drug resistance. As an example, we performed a proliferation experiment using paired baseline and refractory samples of a CLL patient who relapsed on ibrutinib monotherapy. Baseline samples showed strong inhibition of spheroid growth upon *in vitro* ibrutinib treatment whereas spheroid growth of refractory samples was barely affected by *in vitro* ibrutinib treatment (Figure 5A). Notably, refractory cells displayed spheroid growth without any *in vitro* stimulation. The 3D model described here can also be used for more traditional cytotoxic drug screening using either a flow cytometry or live-cell imaging readout. Direct stimulation of CLL cells with IL-2/15/21/CpG resulted in venetoclax resistance similar to 3D coculture with 3T40 fibroblasts, reaching a maximum of 70% specific apoptosis upon 10µM venetoclax (Figure 5B). Stimulation of T cells using αCD3/αCD28 resulted in comparable resistance levels, indicating that T cells are able to confer resistance to CLL cells in this system. This drug resistance was associated with the upregulation of the anti-apoptotic Bcl-2 family members Bcl-XL and Mcl-1 (Figure 5C-D). Aside from proliferation studies, live-cell imaging can also be applied for large-scale drug screening, thereby providing real-time data in addition to endpoint analysis.

**Figure 5.**
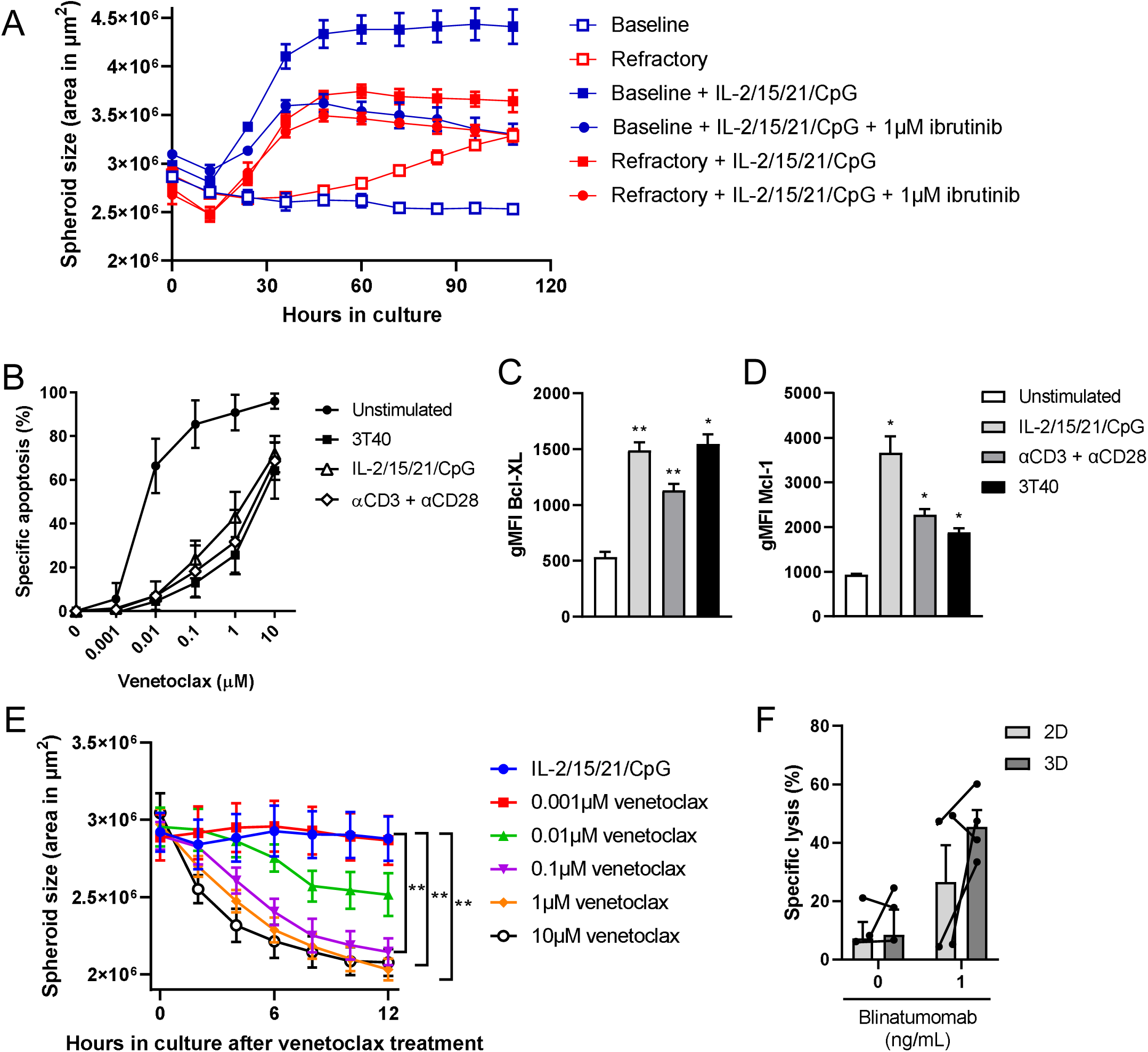
CLL spheroids enable the study of drug resistance and T cell cytotoxicity. **A)** Paired baseline and refractory samples of an ibrutinib-treated CLL patient were stimulated with IL-2/15/21/CpG and cultured in ULA plates (3D) for 4 days in the presence of 1µM ibrutinib. Culture plates were placed in an IncuCyte live-cell imager which imaged and quantified the spheroid area every 12 hours. The data represents 6 spheroids from 1 patient cultured in each condition. Error bars represent the mean ± SD. **B-E)** CLL samples were stimulated as indicated and/or cocultured with 3T40 fibroblast in ULA plates (3D) for 1 week. Afterwards, cells were incubated with a titration of 0.001-10µM venetoclax for an additional 24 hours, during which the culture plates were placed in an IncuCyte live-cell imager which imaged and quantified spheroid area every hour **(E)**. Viability was measured by flow cytometry using DiOC6 and TO-PRO-3 viability dyes **(B)**, including expression of Bcl-XL **(C)** and Mcl-1 **(D)**. Error bars represent the mean ± SEM (n = 3), *p<0.05, **p<0.01 (paired t-test). Each group was compared with the unstimulated control group. **F)** CTV-labeled CLL cells were cocultured with healthy donor T cells in a 4:1 effector to target ratio in the presence of 1ng/mL blinatumomab for 24 hours. Viability was measured by flow cytometry using MitoTracker Orange and TO-PRO-3 viability dyes. Error bars represent the mean ± SEM (n = 4).

Treatment with 0.1-10µM venetoclax significantly decreased spheroid size in a dose-dependent manner (Figure 5E). Furthermore, as proof-of-principle, we performed a T cell cytotoxicity assay via coculture of healthy donor T cells with CLL cells in the presence of the CD3xCD19 bispecific antibody blinatumomab. Coculture in a 4:1 effector to target ratio resulted in specific lysis of CLL cells of 26% in 2D cultures versus 45% in 3D cultures. Notably, 2 healthy donor samples did not respond in standard 2D cultures, whereas they did demonstrate cytotoxicity within the CLL spheroids, thus potentially promoting T cell killing capacity in a 3D conformation (Figure 5F). These data show that conventional T cell cytotoxicity assays are compatible with 3D cultures, thereby opening up new immunotherapies that can be tested in this 3D model, such as antibodies, bispecifics, and CAR T cells.

## Discussion

We developed a 3D culture model using PBMCs from CLL patients, with incorporation of key aspects of CLL LN biology. Based on the 3T40-CLL coculture model^18^, we utilized T cells as a source of physiological CD40L by mimicking the CLL:T cell composition of the CLL LN *in vitro*. By using ULA plates, we created a model that only requires common laboratory equipment and which can thus easily be adopted by other research groups. Using only soluble factors for CLL cell stimulation, the IL-2/15/21/CpG cocktail we presented here can easily be adapted for more specialized applications, such as the αCD3/αCD28 T cell stimulation we also presented. Furthermore, the low required number of cells for seeding and the compatibility with 1 medium exchange per week in the case of long-term cultures makes this model practical to use. Finally, the use of a 96-well plate format enables high-throughput drug screening and still allows accurate comparison with standard 2D culture models.

Our data indicate that enhanced CLL proliferation in 3D cultures is dependent on the presence of T cells. This may be a result of improved interactions of CLL cells with surrounding T cells in a 3D conformation. Noteworthy, although the proliferation index increased in 3D cultures, the maximum number of generations remained the same compared to 2D cultures, suggesting that CLL cells reached a plateau after a certain number of cell divisions. We confirmed that coculture of CLL cells with activated T cells may trigger CLL proliferation^3^, though proliferating CLL cells showed a limited number of cell divisions. A potential explanation is that CD40 stimulation primes CLL cells for proliferation but that additional signals are needed to sustain a proliferative phenotype and drive CLL cells through several cell cycles, such as costimulation with cytokines^3^. Although T cell stimulation does induce the secretion of relevant T cell cytokines, the production and resulting cytokine levels *in vitro* may not be sufficient to sustain CLL proliferation.

Acalabrutinib treatment inhibited spheroid growth and was associated with morphological changes, potentially due to inhibition of Btk-mediated cell adhesion leading to the dissociation of adherent cells. An interesting aspect is how Btk inhibition could affect spheroid growth without the addition of direct BCR stimulation. First, it has been shown that Btk is involved in both TLR-mediated and CD40-mediated CLL proliferation^19-21^. Second, Btk is involved in cell adhesion^22^, and inhibition of Btk may thus affect spheroid growth by disrupting cell-cell interactions within the spheroid. Furthermore, our data demonstrate Bcl-XL and Mcl-1 upregulation by CLL cells after either IL-2/15/21/CpG or T cell stimulation, similar to what is observed in CLL LN samples^18,23^, thereby adding physiological relevance. This could enable ibrutinib-venetoclax synergy studies in an *ex vivo* model. Finally, we observed enhanced CLL proliferation in cocultures with LN-derived stromal cells. The use of CLL-derived or even autologous LN stroma may further support CLL proliferation *in vitro*. Though such material is rare, our data suggest that this 3D model provides a foundation which allows the introduction of additional cell types to the culture and can thus be used for the generation of more complex microenvironments. Moreover, the addition of other cell types, such as follicular dendritic cells or follicular T cells^24,25^ could further influence structure, proliferation, and drug resistance. A distinctive characteristic of CLL LN tissues is the presence of pseudofollicles^26,27^. These are proliferation centers which have not been described in other lymphoproliferative conditions and are enriched in CD40L^+^ T cells and Ki-67^+^ CLL cells compared to surrounding tissue^28^. The emergence of follicle-like structures after prolonged culture in the model could indicate a similar phenotype. Although the presence of follicles can be beneficial by allowing efficient cell-cell communication, this could also result in malnutrition and hypoxia, leading to cell death^29^.

An obvious limitation of the model presented here is the requirement to characterize CLL samples and select them based on CLL:T cell composition. Alternatively, PBMCs can be sorted to acquire the desired CLL:T cell composition. Here, we applied a >10% CD4^+^ T cell cutoff based on the LN biopsy analysis, but it remains unknown where the actual threshold lies in which 3D cultures do not show increased CLL proliferation compared to 2D cultures. Nevertheless, spheroid growth assays can still be carried out independent of CLL:T cell composition, as shown for the ibrutinib-relapse sample that we demonstrated here. A potential oversight of the model is the lack of *in vitro* BCR stimulation using αIgM antibodies, which has been shown to enhance *in vitro* CLL proliferation in combination with costimulatory signals, including a combination of CD40 and cytokine signaling^30^. Though BCR signaling has been acknowledged as a key mechanism for CLL progression *in vivo*^*31*^, we avoided the use of *in vitro* BCR stimulation to exclude differences between IGHV-mutated and IGHV-unmutated CLL samples.

In summary, we present a novel culture system that underlines the role of T cells in CLL proliferation and is capable of sustained growth of CLL cells. In addition to being a scalable, multi-purpose platform, this model may enable the investigation of the outgrowth of genetic (sub)clones^32,33^ and spatial phenotyping of both tumor and non-malignant cells, thereby uncovering new biomarkers and targets for therapy. The model described here permits investigation of CLL cells in the context of their natural protective niche and what role this support plays in CLL drug resistance, thereby opening many new avenues for clinically useful applications.

## Acknowledgements

The authors thank Tineke de Jong and Lisa van Baarsen for providing primary lymph node fibroblasts for the experiment shown in Figure 2F.

This project CLL Lymph node Organoids for Screens and Intelligent Testing (CLOSIT) is in collaboration with Acerta Pharma and Bristol Myers Squibb. The collaboration project is financed by the Ministry of Economic Affairs by means of the PPP Allowance made available by the Top Sector Life Sciences & Health to stimulate public-private partnerships.

## Author contribution statement

MVH designed the study, performed research, analyzed data, and wrote the manuscript. BFvD and EP performed research. DdR processed and characterized patient samples and analyzed data. DL analyzed data and critically reviewed the manuscript. RL acquired financial support and critically reviewed the manuscript. AP provided patient samples and wrote the manuscript. PDM acquired financial support and critically reviewed the manuscript. EE designed the study, acquired financial support and wrote the manuscript.

## Figure Legends

**Supplementary Figure 1. CLL proliferation cocktail optimization. A-C)** CFSE-labeled CLL cells were stimulated as indicated and cultured in 3D spheroid cultures for 1 week. Afterwards, cells were measured using flow cytometry and the percentage of dividing cells was quantified. **(A)** Results of single cytokine or CpG stimulation. **(B)** Results of dual cytokine stimulation or single cytokine stimulation in combination with CpG. **(C)** Results of dual and triple cytokine stimulation in combination with CpG. Bars represent the mean ± SEM (n = 9), **p<0.01, ****p<0.0001 (paired t-test). Each group was compared with the unstimulated control group.

**Supplementary Figure 2. Overview of T cell proliferation data. A-D)** CTV-labeled CLL samples were stimulated as indicated and cultured in standard round-bottom 96 well plates (2D) or ULA plates (3D) for 8 days. Afterwards, cells were measured using flow cytometry, and both the percentage of dividing cells **(A, C)** and the proliferation index were quantified **(B, D, E)** for both CD4^+^ and CD8^+^ T cells. Bars represent the mean ± SEM (n = 10), *p<0.05, **p<0.01, ***p<0.001, ****p<0.0001 (two-way ANOVA). Each group was compared with the unstimulated control group.

## Notes

### Competing Interest Statement

R. Loos is an employee and equity holder of Bristol Myers Squibb.

## References

1. van Attekum MH, Eldering E, Kater AP. Chronic lymphocytic leukemia cells are active participants in microenvironmental cross-talk. Haematologica. 2017;102(9):1469–1476.

2. van Gent R, Kater AP, Otto SA, et al. In vivo dynamics of stable chronic lymphocytic leukemia inversely correlate with somatic hypermutation levels and suggest no major leukemic turnover in bone marrow. Cancer Res. 2008;68(24):10137–10144.

3. Pascutti MF, Jak M, Tromp JM, et al. IL-21 and CD40L signals from autologous T cells can induce antigen-independent proliferation of CLL cells. Blood. 2013;122(17):3010–3019.

4. Simonetti G, Bertilaccio MT, Ghia P, Klein U. Mouse models in the study of chronic lymphocytic leukemia pathogenesis and therapy. Blood. 2014;124(7):1010–1019.

5. Barbaglio F, Belloni D, Scarfo L, et al. Three-dimensional co-culture model of chronic lymphocytic leukemia bone marrow microenvironment predicts patient-specific response to mobilizing agents. Haematologica. 2021;106(9):2334–2344.

6. Svozilova H, Plichta Z, Proks V, et al. RGDS-Modified Superporous Poly(2-Hydroxyethyl Methacrylate)-Based Scaffolds as 3D In Vitro Leukemia Model. Int J Mol Sci. 2021;22(5).

7. Ghia P, Strola G, Granziero L, et al. Chronic lymphocytic leukemia B cells are endowed with the capacity to attract CD4+, CD40L+ T cells by producing CCL22. Eur J Immunol. 2002;32(5):1403–1413.

8. Haselager MV, Kielbassa K, Ter Burg J, et al. Changes in Bcl-2 members after ibrutinib or venetoclax uncover functional hierarchy in determining resistance to venetoclax in CLL. Blood. 2020;136(25):2918–2926.

9. Gupta R, Yan XJ, Barrientos J, et al. Mechanistic Insights into CpG DNA and IL-15 Synergy in Promoting B Cell Chronic Lymphocytic Leukemia Clonal Expansion. J Immunol. 2018;201(5):1570–1585.

10. Decker T, Bogner C, Oelsner M, Peschel C, Ringshausen I. Antiapoptotic effect of interleukin-2 (IL-2) in B-CLL cells with low and high affinity IL-2 receptors. Ann Hematol. 2010;89(11):1125–1132.

11. Decker T, Schneller F, Sparwasser T, et al. Immunostimulatory CpG-oligonucleotides cause proliferation, cytokine production, and an immunogenic phenotype in chronic lymphocytic leukemia B cells. Blood. 2000;95(3):999–1006.

12. de Totero D, Meazza R, Zupo S, et al. Interleukin-21 receptor (IL-21R) is up-regulated by CD40 triggering and mediates proapoptotic signals in chronic lymphocytic leukemia B cells. Blood. 2006;107(9):3708–3715.

13. Griebel P, Beskorwayne T, Van den Broeke A, Ferrari G. CD40 signaling induces B cell responsiveness to multiple members of the gamma chain-common cytokine family. Int Immunol. 1999;11(7):1139–1147.

14. Herishanu Y, Perez-Galan P, Liu D, et al. The lymph node microenvironment promotes B-cell receptor signaling, NF-kappaB activation, and tumor proliferation in chronic lymphocytic leukemia. Blood. 2011;117(2):563–574.

15. Haselager MV, Kater AP, Eldering E. Proliferative Signals in Chronic Lymphocytic Leukemia; What Are We Missing? Front Oncol. 2020;10:592205.

16. Hallaert DY, Jaspers A, van Noesel CJ, van Oers MH, Kater AP, Eldering E. c-Abl kinase inhibitors overcome CD40-mediated drug resistance in CLL: implications for therapeutic targeting of chemoresistant niches. Blood. 2008;112(13):5141–5149.

17. de Weerdt I, Hofland T, de Boer R, et al. Distinct immune composition in lymph node and peripheral blood of CLL patients is reshaped during venetoclax treatment. Blood Adv. 2019;3(17):2642–2652.

18. Tromp JM, Tonino SH, Elias JA, et al. Dichotomy in NF-kappaB signaling and chemoresistance in immunoglobulin variable heavy-chain-mutated versus unmutated CLL cells upon CD40/TLR9 triggering. Oncogene. 2010;29(36):5071–5082.

19. Slinger E, Thijssen R, Kater AP, Eldering E. Targeting antigen-independent proliferation in chronic lymphocytic leukemia through differential kinase inhibition. Leukemia. 2017;31(12):2601–2607.

20. Herman SE, Gordon AL, Hertlein E, et al. Bruton tyrosine kinase represents a promising therapeutic target for treatment of chronic lymphocytic leukemia and is effectively targeted by PCI-32765. Blood. 2011;117(23):6287–6296.

21. Guo A, Lu P, Galanina N, et al. Heightened BTK-dependent cell proliferation in unmutated chronic lymphocytic leukemia confers increased sensitivity to ibrutinib. Oncotarget. 2016;7(4):4598–4610.

22. de Rooij MF, Kuil A, Geest CR, et al. The clinically active BTK inhibitor PCI-32765 targets B-cell receptor- and chemokine-controlled adhesion and migration in chronic lymphocytic leukemia. Blood. 2012;119(11):2590–2594.

23. Smit LA, Hallaert DY, Spijker R, et al. Differential Noxa/Mcl-1 balance in peripheral versus lymph node chronic lymphocytic leukemia cells correlates with survival capacity. Blood. 2007;109(4):1660–1668.

24. Burger JA, Quiroga MP, Hartmann E, et al. High-level expression of the T-cell chemokines CCL3 and CCL4 by chronic lymphocytic leukemia B cells in nurselike cell cocultures and after BCR stimulation. Blood. 2009;113(13):3050–3058.

25. Pedersen IM, Kitada S, Leoni LM, et al. Protection of CLL B cells by a follicular dendritic cell line is dependent on induction of Mcl-1. Blood. 2002;100(5):1795–1801.

26. Herreros B, Rodriguez-Pinilla SM, Pajares R, et al. Proliferation centers in chronic lymphocytic leukemia: the niche where NF-kappaB activation takes place. Leukemia. 2010;24(4):872–876.

27. Ponzoni M, Doglioni C, Caligaris-Cappio F. Chronic lymphocytic leukemia: the pathologist’s view of lymph node microenvironment. Semin Diagn Pathol. 2011;28(2):161–166.

28. Vandewoestyne ML, Pede VC, Lambein KY, et al. Laser microdissection for the assessment of the clonal relationship between chronic lymphocytic leukemia/small lymphocytic lymphoma and proliferating B cells within lymph node pseudofollicles. Leukemia. 2011;25(5):883–888.

29. Verjans ET, Doijen J, Luyten W, Landuyt B, Schoofs L. Three-dimensional cell culture models for anticancer drug screening: Worth the effort? J Cell Physiol. 2018;233(4):2993–3003.

30. Schleiss C, Ilias W, Tahar O, et al. BCR-associated factors driving chronic lymphocytic leukemia cells proliferation ex vivo. Sci Rep. 2019;9(1):701.

31. Burger JA, Chiorazzi N. B cell receptor signaling in chronic lymphocytic leukemia. Trends Immunol. 2013;34(12):592–601.

32. Gutierrez C, Wu CJ. Clonal dynamics in chronic lymphocytic leukemia. Blood Adv. 2019;3(22):3759–3769.

33. Landau DA, Carter SL, Stojanov P, et al. Evolution and impact of subclonal mutations in chronic lymphocytic leukemia. Cell. 2013;152(4):714–726.

