## Supplementary figures and images for "*In vitro* lymph node-mimicking 3D model displays long-term T cell-dependent CLL proliferation and survival"

### Supplemental Figure 1

#### Slide 1
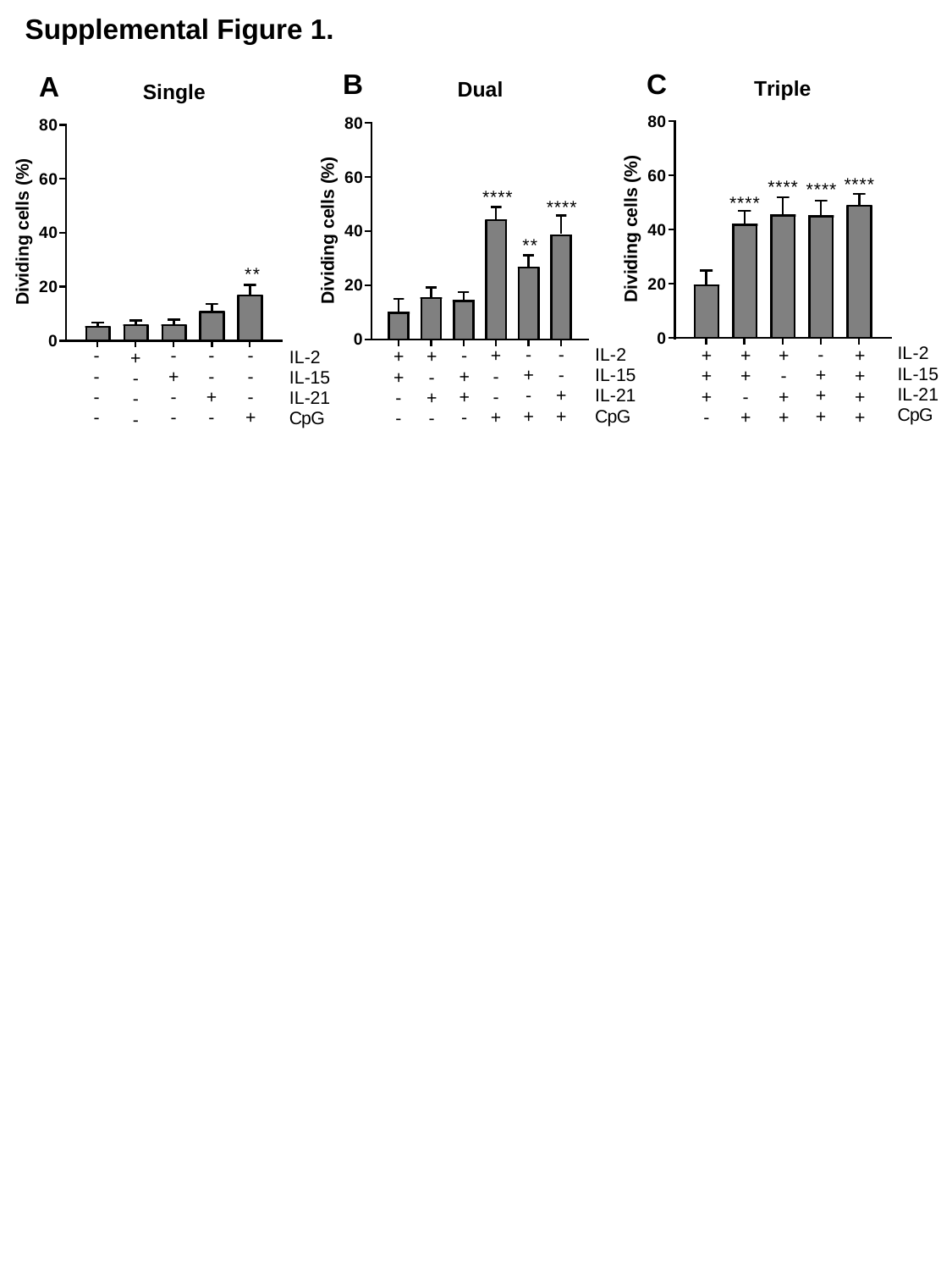

Supplemental Figure 1.
B
C
A

### Supplemental Figure 2

#### Slide 1
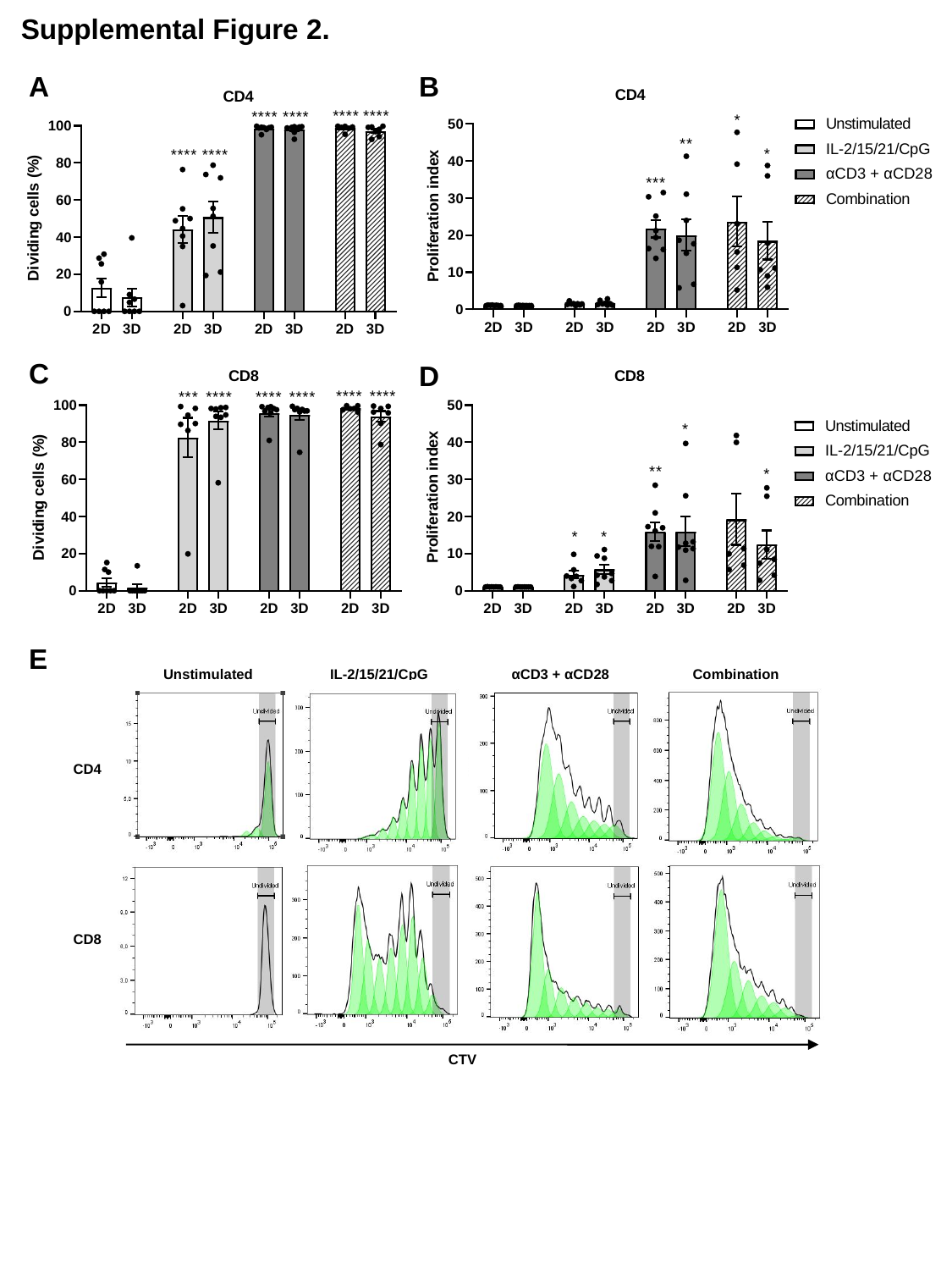

Supplemental Figure 2.
A
B
C
D
E
Unstimulated
IL-2/15/21/CpG
αCD3 + αCD28
Combination
CD4
CD8
CTV
