## Supplemental Table 1 for "*In vitro* lymph node-mimicking 3D model displays long-term T cell-dependent CLL proliferation and survival"

| sample ID | experiment | sample date | age | gender | IgHV mutation status | treatment | article figure |
|---|---|---|---|---|---|---|---|
| H1580001T1 | Biobank (PB) | 30-12-2020 | 74 | M | unknown | none | 1A-B |
| H1580001T1 | Biobank (LN) | 30-12-2020 | 74 | M | unknown | none | 1A-B |
| H1580002T1 | Biobank (PB) | 4-1-2021 | 52 | M | unknown | none | 1A-B |
| H1580002T1 | Biobank (LN) | 4-1-2021 | 52 | M | unknown | none | 1A-B |
| H1580004T1 | Biobank (PB) | 18-1-2021 | 59 | M | unknown | none | 1A-B |
| H1580004T1 | Biobank (LN) | 18-1-2021 | 59 | M | unknown | none | 1A-B |
| H1580005T1 | Biobank (PB) | 19-1-2021 | 72 | M | unknown | none | 1A-B |
| H1580005T1 | Biobank (LN) | 19-1-2021 | 72 | M | unknown | none | 1A-B |
| H1580006T1 | Biobank (PB) | 19-1-2021 | 74 | F | unknown | none | 1A-B |
| H1580006T1 | Biobank (LN) | 19-1-2021 | 74 | F | unknown | none | 1A-B |
| H1580009T1 | Biobank (PB) | 27-1-2021 | 60 | M | unknown | none | 1A-B |
| H1580009T1 | Biobank (LN) | 27-1-2021 | 60 | M | unknown | none | 1A-B |
| H1580010T1 | Biobank (PB) | 3-2-2021 | 64 | M | unknown | none | 1A-B |
| H1580010T1 | Biobank (LN) | 3-2-2021 | 64 | M | unknown | none | 1A-B |
| H1580011T1 | Biobank (PB) | 8-2-2021 | 70 | M | unknown | none | 1A-B |
| H1580011T1 | Biobank (LN) | 8-2-2021 | 70 | M | unknown | none | 1A-B |
| H1580016T1 | Biobank (PB) | 2-3-2021 | 82 | M | unknown | none | 1A-B |
| H1580016T1 | Biobank (LN) | 2-3-2021 | 82 | M | unknown | none | 1A-B |
| H1580018T1 | Biobank (PB) | 23-2-2021 | 71 | M | unknown | none | 1A-B |
| H1580018T1 | Biobank (LN) | 23-2-2021 | 71 | M | unknown | none | 1A-B |
| H1580019T1 | Biobank (PB) | 24-2-2021 | 35 | M | unknown | none | 1A-B |
| H1580019T1 | Biobank (LN) | 24-2-2021 | 35 | M | unknown | none | 1A-B |
| H1580021T1 | Biobank (PB) | 25-2-2021 | 66 | M | unknown | none | 1A-B |
| H1580021T1 | Biobank (LN) | 25-2-2021 | 66 | M | unknown | none | 1A-B |
| H1580023T1 | Biobank (PB) | 3-3-2021 | 65 | M | unknown | none | 1A-B |
| H1580023T1 | Biobank (LN) | 3-3-2021 | 65 | M | unknown | none | 1A-B |
| H1580025T1 | Biobank (PB) | 2-3-2021 | 68 | M | unknown | none | 1A-B |
| H1580025T1 | Biobank (LN) | 2-3-2021 | 68 | M | unknown | none | 1A-B |
| H1580028T1 | Biobank (PB) | 16-3-2021 | 60 | F | unknown | none | 1A-B |
| H1580028T1 | Biobank (LN) | 16-3-2021 | 60 | F | unknown | none | 1A-B |
| H1580029T1 | Biobank (PB) | 9-3-2021 | 67 | F | unknown | none | 1A-B |
| H1580029T1 | Biobank (LN) | 9-3-2021 | 67 | F | unknown | none | 1A-B |
| H1580030T1 | Biobank (PB) | 15-3-2021 | 66 | F | unknown | none | 1A-B |
| H1580030T1 | Biobank (LN) | 15-3-2021 | 66 | F | unknown | none | 1A-B |
| H1580031T1 | Biobank (PB) | 23-3-2021 | 68 | M | unknown | none | 1A-B |
| H1580031T1 | Biobank (LN) | 23-3-2021 | 68 | M | unknown | none | 1A-B |
| H1580032T1 | Biobank (PB) | 15-3-2021 | 86 | M | unknown | none | 1A-B |
| H1580032T1 | Biobank (LN) | 15-3-2021 | 86 | M | unknown | none | 1A-B |
| H1580033T1 | Biobank (PB) | 16-3-2021 | 56 | F | unknown | none | 1A-B |
| H1580033T1 | Biobank (LN) | 16-3-2021 | 56 | F | unknown | none | 1A-B |
| H1580034T1 | Biobank (PB) | 17-3-2021 | 58 | M | unknown | none | 1A-B |
| H1580034T1 | Biobank (LN) | 17-3-2021 | 58 | M | unknown | none | 1A-B |
| H1580036T1 | Biobank (PB) | 24-3-2021 | 75 | M | unknown | none | 1A-B |
| H1580036T1 | Biobank (LN) | 24-3-2021 | 75 | M | unknown | none | 1A-B |
| H1580038T1 | Biobank (PB) | 30-3-2021 | 73 | M | unknown | none | 1A-B |
| H1580038T1 | Biobank (LN) | 30-3-2021 | 73 | M | unknown | none | 1A-B |
| H1580039T1 | Biobank (PB) | 30-3-2021 | 63 | F | unknown | none | 1A-B |
| H1580039T1 | Biobank (LN) | 30-3-2021 | 63 | F | unknown | none | 1A-B |
| 2967 | Confocal staining | 6-4-2020 | 69 | F | unknown | none | 1C |
| 2367 | Immunophenotyping | 18-9-2018 | 68 | F | unknown | none | 1D |
| 2549 | Immunophenotyping | 23-4-2019 | 77 | M | unknown | none | 1D |
| 3067 | Immunophenotyping | 26-6-2020 | 60 | F | unknown | none | 1D |
| 2218 | Immunophenotyping | 15-5-2018 | 51 | M | unknown | none | 1D |
| 2973 | Immunophenotyping | 14-4-2020 | 70 | M | unknown | none | 1D |
| 3065 | Immunophenotyping | 24-6-2020 | 62 | F | unknown | none | 1D |
| 3236 | Immunophenotyping | 26-10-2020 | 70 | F | unknown | none | 1D |
| 2244 | Immunophenotyping | 5-6-2018 | 49 | M | unknown | none | 1D |
| 1093 | CLL/T cell activation/proliferation | 5-9-2011 | 53 | M | mutated | none | 2A-B, 2C, 2E, 2G, S2A-D |
| 1448 | CLL/T cell activation/proliferation | 6-2-2014 | 63 | M | mutated | none | 2A-B, 2C, 2E, 2G, S2A-D |
| 1093 | CLL/T cell activation/proliferation | 5-9-2011 | 53 | M | mutated | none | 2A-B, 2C, 2E, 2G, S2A-D |
| 1201 | CLL/T cell activation/proliferation | 11-5-2012 | 61 | F | unknown | none | 2A-B, 2C, 2E, 2G, S2A-D |
| 1093 | CLL/T cell activation/proliferation | 5-9-2011 | 53 | M | mutated | none | 2A-E, 2G, S2A-D |
| 1201 | CLL/T cell activation/proliferation | 11-5-2012 | 61 | F | unknown | none | 2A-B, 2C, 2E, 2G, S2A-D |
| 1201 | CLL/T cell activation/proliferation | 11-5-2012 | 61 | F | unknown | none | 2A-B, 2C, 2E, 2G, S2A-D |
| 1239 | CLL/T cell activation/proliferation | 24-7-2012 | 71 | F | unmutated | none | 2A-B, 2C, 2E, 2G, S2A-E |
| 1201 | CLL/T cell activation/proliferation | 11-5-2012 | 61 | F | unknown | none | 2A-B, 2C, 2E, 2G, S2A-D |
| 1239 | CLL/T cell activation/proliferation | 24-7-2012 | 71 | F | unmutated | none | 2A-B, 2C, 2E, 2G, S2A-D |
| 187 | LN stroma proliferation | 15-11-2005 | 58 | F | mutated | none | 2F |

| | | | | | | | |
|---|---|---|---|---|---|---|---|
| 197 | LN stroma proliferation | 15-12-2005 | 60 | M | unmutated | 2x chloorambucil, radiotherapy, fludarabine | 2F |
| 1673 | LN stroma proliferation | 3-8-2015 | 40 | M | unmutated | none | 2F |
| 333 | Long-term proliferation cultures | 24-5-2002 | 86 | F | unmutated | none | 3A-B |
| 1201 | Long-term proliferation cultures | 11-5-2012 | 61 | F | unknown | none | 3A, 3C |
| 2089 | Long-term proliferation cultures | 16-1-2018 | 65 | F | unknown | none | 3A, 3D |
| 1448 | Long-term proliferation cultures | 6-2-2014 | 63 | M | mutated | none | 3A |
| 361 | Long-term proliferation cultures | 29-8-2000 | 69 | M | unmutated | chloorambucil, 2x fludarabine, CHOP+R, radiotherapy | 3A |
| 2089 | Long-term proliferation cultures | 16-1-2018 | 65 | F | unknown | none | 3A |
| 273 | Ibrutinib proliferation assay | 29-6-2006 | 77 | M | mutated | none | 4A |
| 1065 | Ibrutinib proliferation assay | 8-6-2011 | 73 | M | mutated | none | 4A-B |
| 1065 | Ibrutinib proliferation assay | 8-6-2011 | 73 | M | mutated | none | 4A-C |
| 2791 | Ibrutinib proliferation assay | 11-12-2019 | 55 | M | unknown | none | 4B |
| 3194 | Ibrutinib proliferation assay | 21-9-2020 | 62 | M | mutated | FCR | 4B |
| 222 | Ibrutinib proliferation assay | 28-2-2006 | 54 | M | polyclonal | none | 4B |
| 273 | Cohesion analysis | 29-6-2006 | 77 | M | mutated | none | 4E |
| 1065 | Cohesion analysis | 8-6-2011 | 73 | M | mutated | none | 4E |
| 2791 | Cohesion analysis | 11-12-2019 | 55 | M | unknown | none | 4D-E |
| 1546 | Refractory ibrutinib sample | 24-9-2014 | 50 | M | unmutated | FCR | 5A |
| 3104 | Refractory ibrutinib sample | 21-7-2020 | 56 | M | unmutated | FCR, ibrutinib | 5A |
| 197 | Venetoclax sensitivity | 15-12-2005 | 60 | M | unmutated | 2x chloorambucil, radiotherapy, fludarabine | 5B |
| 1557 | Venetoclax sensitivity | 17-10-2014 | 64 | F | mutated | none | 5B |
| 1873 | Venetoclax sensitivity | 16-1-2017 | 54 | F | mutated | none | 5B |
| 64 | Intracellular Bcl-2 FACS | 16-12-2004 | 66 | F | mutated | none | 5C-D |
| 1085 | Intracellular Bcl-2 FACS | 9-8-2011 | 70 | M | mutated | none | 5C-D |
| 2029 | Intracellular Bcl-2 FACS | 12-9-2017 | 77 | F | mutated | none | 5C-D |
| 702 | 3D cytotoxicity | 16-10-2008 | 64 | F | mutated | none | 5E |
| 721 | 3D cytotoxicity | 18-12-2008 | 85 | M | mutated | none | 5E |
| 708 | 3D cytotoxicity | 13-11-2008 | 68 | F | mutated | none | 5E |
| 839 | 3D cytotoxicity | 9-9-2009 | 71 | F | mutated | none | 5E |
| 123 | T cell cytotoxicity | 25-7-2003 | 61 | M | mutated | none | 5F |
| 1659 | T cell cytotoxicity | 25-6-2015 | 57 | M | mutated | none | 5F |
| HD 93 | T cell cytotoxicity | 1-12-2021 | 63 | M | - | - | 5F |
| HD 96 | T cell cytotoxicity | 1-12-2021 | 62 | M | - | - | 5F |
| 1753 | 2D-3D proliferation optimization | 21-3-2016 | 62 | F | unmutated | FCR, chloorambucil | S1A-C |
| 1797 | 2D-3D proliferation optimization | 30-8-2016 | 74 | F | mutated | none | S1A-C |
| 1439 | 2D-3D proliferation optimization | 13-1-2014 | 57 | M | mutated | none | S1A-C |
| 1673 | 2D-3D proliferation optimization | 3-8-2015 | 40 | M | unmutated | none | S1A-C |
| 1808 | 2D-3D proliferation optimization | 12-9-2016 | 46 | M | unmutated | none | S1A-C |
| 1254 | 2D-3D proliferation optimization | 11-9-2012 | 62 | F | mutated | none | S1A-C |
| 1254 | 2D-3D proliferation optimization | 11-9-2012 | 62 | F | mutated | none | S1A-C |
| 1857 | 2D-3D proliferation optimization | 2-1-2017 | 84 | M | unmutated | none | S1A-C |
| 2038 | 2D-3D proliferation optimization | 2-10-2017 | 68 | F | mutated | none | S1A-C |
