## Supplemental information for "*In vitro* lymph node-mimicking 3D model displays long-term T cell-dependent CLL proliferation and survival"

**Protocol 1: 3D cell culture.**

| **Resource** | **Article number** | **Supplier** |
| --- | --- | --- |
| PBMCs | n/a | n/a |
| Iscove’s Modified Dulbecco’s Medium | 12440053 | Gibco (Waltham, MA, USA) |
| Fetal bovine serum | 10437028 | Gibco (Waltham, MA, USA) |
| Penillin/streptomycin | 15140-22 | Invitrogen (Waltham, MA, USA) |
| 96-well clear round-bottom ultra-low attachment plates | 7007 | Corning (Corning, NY, USA) |
| Recombinant human IL-2 | 200-02-B | Peprotech (London, United Kingdom) |
| Recombinant human IL-15 | 200-15-B | Peprotech (London, United Kingdom) |
| Recombinant human IL-21 | PHC0215 | Thermo Scientific (Waltham, MA, USA) |
| ODN 2006 (CpG) | tlrl-2006 | Invivogen (San Diego, CA, USA) |
| αCD3 (clone 1XE) | n/a | of your choice |
| αCD28 (clone 15E8) | n/a | of your choice |
| CellTrace™ Violet Cell Proliferation Kit, for flow cytometry | C34557 | Thermo Scientific (Waltham, MA, USA) |
| Drugs | n/a | of your choice |
| Pipettes and corresponding plastic-ware | n/a | of your choice |
| Humidified incubator at 37⁰C and 5% CO_2_ | n/a | of your choice |
| Water bath with controlled temperature | n/a | of your choice |
| Centrifuge | n/a | of your choice |
| Sterile biosafety cabinet | n/a | of your choice |

1. Select CLL-derived PB sample with >10% CD3^+^CD4^+^ T cells (guideline: 10-20% CD3^+^CD4^+^ T cells and 50-80% CD5^+^CD19^+^ CLL cells) and retrieve ampule from liquid nitrogen storage.

2. Thaw CLL-derived PB sample.

a. Thaw ampule by swinging the ampule in a water bath at 37°C until the ice is dissolved.

b. Under sterile conditions, in a laminar flow hood, transfer 2mL of cell suspension into a 50mL tube.

c. Rinse the ampule with 2mL cold thawing medium (IMDM, 20% FCS, 1% pen/strep, 4°C) and add drop-wise to the 50mL tube.

d. Add 20mL of cold thawing medium drop-wise to the 50mL tube.

e. Incubate cells for 20 minutes at room temperature in the dark.

f. Centrifuge cells for 5 minutes at 578 x g.

g. Discard supernatant.

h. Resuspend the cells in 10mL warm culture medium (IMDM, 10% FCS, 1% pen/strep, 37°C).

i. Count the cells.

j. Dilute cell suspension to a minimum of 1 x 10^5^ cells/mL (1 x 10^4^ cells/spheroid) and a maximum of 3 x 10^6^ cells/mL (3 x 10^5^ cells/spheroid).

3. Optional: Label the CLL-derived PBMCs with CellTrace Violet.

a. Transfer 5 x 10^6^ cells in 15mL tube

b. Add 5mL PBS to the 15mL tube.

c. Centrifuge cells for 5 minutes at 578 x g.

d. Discard supernatant.

e. Resuspend cells in 1mL 5µM CellTrace Violet solution.

f. Incubate the cells for 15 minutes in a water bath at 37⁰C in the dark while shaking.

g. Add 5mL medium to the 15mL tube.

h. Centrifuge cells for 5 minutes at 578 x g.

i. Resuspend the cells in medium to a concentration of 8 x 10^5^ cells/mL.

4. Prepare 3x concentrated B cell stimulation cocktail in medium (IMDM, 10% FCS, 1% pen/strep).

a. 75ng/mL IL-2 (25ng/mL final concentration).

b. 75ng/mL IL-15 (25ng/mL final concentration).

c. 75ng/mL IL-21 (25ng/mL final concentration).

d. 3µg/mL CpG (1µg/mL final concentration).

5. Optional: Prepare 3x concentrated T cell stimulation cocktail in medium (IMDM, 10% FCS, 1% pen/strep).

a. 273ng/mL anti-CD3 (clone 1XE) (91ng/mL final concentration).

b. 9µg/mL anti-CD28 (clone 15E8) (3µg/mL final concentration).

c. Either substitute for or combine with B cell stimulation cocktail depending on experimental aim.

6. Optional: Prepare cell suspension with stromal or myeloid cells in a concentration of minimum 2 x 10^3^ cells/mL and maximum 6 x 10^5^ cells/mL, taking into account a 1:5 ratio of stroma:PBMC cells (for example: 6 x 10^4^ stromal cells per 3 x 10^5^ PBMCs).

7. Optional: Prepare a 3x concentrated drug solution.

8. Setup 3D cultures.

a. Pipette 100µL cell suspension per well in a 96-well ultra-low attachment plate (1 x 10^4^ - 3 x 10^5^ cells/spheroid).

b. Add 100µL 3x concentrated B cell stimulation cocktail solution or medium per well.

c. Add 100µL medium per well for a total volume of 300µL per well (optionally: substitute with 100µL stromal/myeloid cell suspension or 100µL 3x concentrated drug solution).

d. Centrifuge ultra-low attachment plate for 10 minutes at 200 x g.

e. Place ultra-low attachment plate in humidified incubator (37⁰C, 5% CO_2_) and incubate overnight to allow spheroid formation.

9. Culture spheroids for 24 hours up to 7 weeks depending on experimental aim, downstream workflow and readout (guideline: cell activation: 3-5 days; proliferation assays: 3-7 days; drug screening: 24 h - 1 week; long-term (proliferation) assays: up to approximately 7 weeks while refreshing medium every week). Specific protocols for each downstream workflow are provided below.

10. Optional: Refresh medium weekly when culturing cells for longer than 1 week.

a. Slowly remove the supernatant using a pipette without touching the spheroid to minimize effects on spheroid structure.

b. Prepare fresh medium supplemented with relevant stimuli.

c. Slowly add fresh supplemented medium via the side of the well to minimize effects on spheroid structure.

**Protocol 2: Flow cytometry.**

| **Resource** | **Article number** | **Supplier** |
| --- | --- | --- |
| Pipettes and corresponding plastic-ware | n/a | of your choice |
| 96-well clear round-bottom plates | 3799 | Corning (Corning, NY, USA) |
| Centrifuge | n/a | of your choice |
| Plate shaker | n/a | of your choice |
| Fetal bovine serum | 10437028 | Gibco (Waltham, MA, USA) |
| Phosphate-buffered saline | n/a | Fresenius Kahi (Bad Homburg vor der Höhe, Germany) |
| BD Cytofix/Cytoperm™ | 554772 | BD Biosciences (Franklin Lakes, NJ, USA) |
| BD Perm/Wash™ | 554723 | BD Biosciences (Franklin Lakes, NJ, USA) |
| Antibodies | n/a | of your choice |
| Viability dye | n/a | of your choice |
| Flow cytometer | n/a | of your choice |

1. After culture, resuspend the cells to dissociate the spheroids in order to ensure proper flow cytometry staining.

2. Optional step to keep the unused wells of the ultra-low attachment plate sterile: Transfer cells to a non-sterile round-bottom 96-well culture plate.

3. Perform flow cytometry staining.

a. Centrifuge cells for 5 minutes at 578 x g.

b. Wash cells with 100µL 0.5% BSA diluted in PBS.

c. Resuspend cells in 20µL antibody staining.

d. Incubate cells for 30 minutes at 4⁰C on a plate shaker at 450 rpm in the dark.

e. Wash cells 2x with 100µL 0.5% BSA diluted in PBS.

f. Resuspend cells in 100µL 0.5% BSA diluted in PBS.

4. Optional: Perform intracellular flow cytometry staining.

a. Fix and permeabilize cells in 100µL BD Cytofix/Cytoperm solution.

b. Incubate cells for 15 minutes at 4⁰C on a plate shaker at 450 rpm in the dark.

c. Wash cells 4x with 1x BD Perm/Wash solution.

d. Resuspend cells in 20µL antibody staining.

e. Incubate cells for 30 minutes at 4⁰C on a plate shaker at 450 rpm in the dark.

f. Wash cells 2x with 100µL BD Perm/Wash solution.

g. Resuspend cells in 100µL 0.5% BSA diluted in PBS.

5. Measure cells on a flow cytometer.

**Protocol 3: Confocal microscopy.**

| **Resource** | **Article number** | **Supplier** |
| --- | --- | --- |
| Pipettes and corresponding plastic-ware | n/a | of your choice |
| 96-well clear flat-bottom plates | 3596 | Corning (Corning, NY, USA) |
| Centrifuge | n/a | of your choice |
| Plate shaker | n/a | of your choice |
| Agarose | A9539 | Sigma Aldrich (Saint Louis, MO, USA) |
| Microwave | n/a | of your choice |
| Paraformaldehyde | 28906 | Thermo Scientific (Waltham, MA, USA) |
| Triton X-100 | T8787 | Sigma Aldrich (Saint Louis, MO, USA) |
| Fetal bovine serum | 10437028 | Gibco (Waltham, MA, USA) |
| Phosphate-buffered saline | n/a | Fresenius Kahi (Bad Homburg vor der Höhe, Germany) |
| DAPI | D9542 | Sigma Aldrich (Saint Louis, MO, USA) |
| Antibodies | n/a | of your choice |
| µ-Slide 18 well Glass Bottom | 81817 | Ibidi (Gräfelfing, Germany) |
| Confocal microscope | n/a | of your choice |

1. After culture, embed spheroids in 1% agarose.

a. Slowly remove the supernatant using a pipette without touching the spheroid to minimize effects on spheroid structure.

b. Slowly add 200µL 37⁰C 1% agarose solution via the side of the well to minimize effects on spheroid structure.

c. Incubate plate for approximately 10 minutes at room temperature until agarose gels solidify.

2. Perform cell fixation.

a. Insert the small end of the plastic spatula all the way into the well and scoop out the entire agarose gel.

b. Transfer agarose gels into 4% paraformaldehyde solution.

c. Incubate for 10 minutes at room temperature on a plate shaker at 450 rpm.

d. Wash spheroids 3x with PBS.

e. Agarose-embedded spheroids can be stored in PBS at 4⁰C after this step.

3. Optional: Perform cell permeabilization.

a. Transfer agarose gels into 0.1% Triton X-100 solution.

b. Incubate for 10 minutes at room temperature on a plate shaker at 450 rpm.

c. Remove Triton X-100 and wash spheroids 3x with PBS.

4. Block the cells in order to minimize aspecific antibody staining.

a. Transfer agarose gels into 0.5% BSA diluted in PBS for 1 hour at room temperature on a plate shaker at 450rpm.

b. Wash spheroids with PBS.

5. Perform primary antibody staining using directly labeled antibodies or unconjugated antibodies.

a. Per spheroid: pipette 100µL primary antibody solution in a flat-bottom 96 well plate.

b. Transfer spheroids into primary antibody solution.

c. Incubate spheroids overnight at 4⁰C on a plate shaker at 450 rpm in the dark.

d. Per spheroid: Pipette 3 wells with 100µL PBS in a flat-bottom 96 well plate.

e. Wash spheroids 3x with PBS by transferring them from well to well while incubating them for 5 minutes in between each washing step on a plate shaker at 450 rpm in the dark.

6. Optional in the case of unconjugated primary antibodies: Perform secondary antibody staining.

a. Per spheroid: pipette 100µL secondary antibody solution in a flat-bottom 96 well plate.

b. Transfer spheroids into secondary antibody solution.

c. Incubate spheroids for 1 hour at room temperature on a plate shaker at 450 rpm in the dark.

d. Per spheroid: Pipette 3 wells with 100µL PBS in a flat-bottom 96 well plate.

e. Wash spheroids 3x with PBS by transferring them from well to well while incubating them for 5 minutes in between each washing step on a plate shaker at 450 rpm in the dark.

7. Optional in the case of larger antibody panels: Perform tertiary antibody staining using directly labeled antibodies.

a. Per spheroid: pipette 100µL tertiary antibody solution in a flat-bottom 96 well plate.

b. Transfer spheroids into tertiary antibody solution.

c. Incubate spheroids overnight at 4⁰C on a plate shaker at 450 rpm in the dark.

d. Per spheroid: Pipette 3 wells with 100µL PBS in a flat-bottom 96 well plate.

e. Wash spheroids 3x with PBS by transferring them from well to well while incubating them for 5 minutes in between each washing step on a plate shaker at 450 rpm in the dark.

8. Optional: Perform DAPI staining.

a. Per spheroid: pipette 100µL 1x DAPI solution in a flat-bottom 96 well plate.

b. Transfer spheroids into DAPI solution.

c. Incubate spheroids for 30 minutes at room temperature on a plate shaker at 450 rpm in the dark.

d. Per spheroid: Pipette 2 wells with 100µL PBS in a flat-bottom 96 well plate.

e. Wash spheroids with PBS by transferring them from well to well while incubating them for 5 minutes in between each washing step on a plate shaker at 450 rpm in the dark.

9. Analyze agarose-embedded spheroids by confocal microscopy.

a. Transfer agarose-embedded spheroids from PBS onto an 18-well glass-bottom microscopic slide.

b. Push each gel into a well until the spheroid touches the glass surface of the slide.

c. Remove excess agarose.

d. For each spheroid: fill the well with PBS as mounting medium.

e. Analyze by confocal microscopy.

10. Optional for long-term storage: Store the agarose-embedded spheroids at 4⁰C in the dark in PBS. Loss of moisture will dry out and destroy the agarose gels.

**Protocol 4: Cytotoxic drug screen.**

| **Resource** | **Article number** | **Supplier** |
| --- | --- | --- |
| Pipettes and corresponding plastic-ware | n/a | of your choice |
| 96-well clear round-bottom plates | 3799 | Corning (Corning, NY, USA) |
| Centrifuge | n/a | of your choice |
| Plate shaker | n/a | of your choice |
| Fetal bovine serum | 10437028 | Gibco (Waltham, MA, USA) |
| Phosphate-buffered saline | n/a | Fresenius Kahi (Bad Homburg vor der Höhe, Germany) |
| DioC6(3) | D273 | Thermo Scientific (Waltham, MA, USA) |
| TO-PRO-3 iodide (642/661) | T3605 | Thermo Scientific (Waltham, MA, USA) |
| Cytotoxic drugs | n/a | of your choice |
| Antibodies | n/a | of your choice |
| IncuCyte live-cell analysis system | n/a | Sartorius (Goettingen, Germany) |
| Flow cytometer | n/a | of your choice |

1. After culture, perform in vitro drug treatment.

a. Slowly remove the supernatant using a pipette without touching the spheroid to minimize effects on spheroid structure.

b. Prepare 1x concentrated drug solution.

c. Slowly add 200µL 1x concentrated drug solution via the side of the well to minimize effects on spheroid structure.

2. Optional: Perform real-time live-cell imaging analysis.

a. Place ultra-low attachment plate in an IncuCyte live-cell analysis system inside humidified incubator (37⁰C, 5% CO_2_).

b. Set up a scan with the Spheroid, Single, Phase + Brightfield settings using a 10x objective.

c. Schedule scans in an interval of 6 hours.

d. Incubate spheroids for your designated incubation time (guideline: 24 hours up to 1 week).

e. Afterwards, quantify spheroid area using IncuCyte software via the Spheroid analysis.

3. Optional: Perform endpoint flow cytometry analysis.

a. After culture, resuspend the cells to dissociate the spheroids in order to ensure proper flow cytometry staining.

b. Transfer 100µL cell suspension per well to a non-sterile 96-well plate.

c. Add 10µL 0.1µM DioC6 solution per well.

d. Incubate the cells for 20 min at 37⁰C in the dark.

e. Optional: Perform flow cytometry surface staining.

i. Centrifuge cells for 5 minutes at 578 x g.

ii. Wash cells with 100µL 0.5% BSA diluted in PBS.

iii. Resuspend cells in 20µL antibody staining.

iv. Incubate cells for 30 minutes at 4⁰C on a plate shaker at 450 rpm in the dark.

v. Wash cells 2x with 100µL 0.5% BSA diluted in PBS

vi. Resuspend cells in 100µL 0.5% BSA diluted in PBS.

f. Add 5µL 0.2µM TO-PRO-3 solution per well.

g. Incubate the cells for 5 min at 4⁰C in the dark.

h. Measure cells on a flow cytometer.

**Protocol 5: T cell cytotoxicity assay.**

| **Resource** | **Article number** | **Supplier** |
| --- | --- | --- |
| CLL-derived PBMCs | n/a | n/a |
| HD-derived PBMCs | n/a | n/a |
| Pipettes and corresponding plastic-ware | n/a | of your choice |
| 96-well clear round-bottom ultra-low attachment plates | 7007 | Corning (Corning, NY, USA) |
| Iscove’s Modified Dulbecco’s Medium | 12440053 | Gibco (Waltham, MA, USA) |
| Fetal bovine serum | 10437028 | Gibco (Waltham, MA, USA) |
| Penillin/streptomycin | 15140-22 | Invitrogen (Waltham, MA, USA) |
| Phosphate-buffered saline | n/a | Fresenius Kahi (Bad Homburg vor der Höhe, Germany) |
| CellTrace™ Violet Cell Proliferation Kit, for flow cytometry | C34557 | Thermo Scientific (Waltham, MA, USA) |
| DioC6(3) | D273 | Thermo Scientific (Waltham, MA, USA) |
| TO-PRO-3 iodide (642/661) | T3605 | Thermo Scientific (Waltham, MA, USA) |
| Blinatumomab | n/a | Amgen (Thousand Oaks, CA, USA) |
| Venetoclax | A0776 | LKT Laboratories (St Paul, MN, USA) |
| Antibodies | n/a | of your choice |
| Humidified incubator at 37⁰C and 5% CO_2_ | n/a | of your choice |
| Water bath with controlled temperature | n/a | of your choice |
| Centrifuge | n/a | of your choice |
| Sterile biosafety cabinet | n/a | of your choice |
| Plate shaker | n/a | of your choice |
| Flow cytometer | n/a | of your choice |

1. Select CLL-derived PB sample with >10% CD3^+^CD4^+^ T cells (guideline: 10-20% CD3^+^CD4^+^ T cells and 50-80% CD5^+^CD19^+^ CLL cells) and retrieve ampule from liquid nitrogen storage.

2. Thaw CLL-derived PB sample and HD-derived PB sample.

a. Thaw ampule by swinging the ampule in a water bath at 37°C until the ice is dissolved.

b. Under sterile conditions, in a laminar flow hood, transfer 2mL of cell suspension into a 50mL tube.

c. Rinse the ampule with 2mL cold thawing medium (IMDM, 20% FCS, 1% pen/strep, 4°C) and add drop-wise to the 50mL tube.

d. Add 20mL of cold thawing medium drop-wise to the 50mL tube.

e. Incubate cells for 20 minutes at room temperature in the dark.

f. Centrifuge cells for 5 minutes at 578 x g.

g. Discard supernatant.

h. Resuspend the cells in 10mL warm culture medium (IMDM, 10% FCS, 1% pen/strep, 37°C).

i. Count the cells.

3. Label the CLL-derived PBMCs with CellTrace Violet.

a. Transfer 5 x 10^6^ cells in 15mL tube

b. Add 5mL PBS to the 15mL tube.

c. Centrifuge cells for 5 minutes at 578 x g.

d. Discard supernatant.

e. Resuspend cells in 1mL 5µM CellTrace Violet solution.

f. Incubate the cells for 15 minutes in a water bath at 37⁰C in the dark while shaking.

g. Add 5mL medium to the 15mL tube.

h. Centrifuge cells for 5 minutes at 578 x g.

i. Resuspend the cells in medium to a concentration of 8 x 10^5^ cells/mL.

4. Set up the T cell killing experiment.

a. Add 50µL CLL-derived PBMCs per well to a sterile 96-well ultra-low attachment plate (4 x 10^4^ target cells per well).

b. Dilute HD-derived PBMCs to a concentration of 3,2 x 10^6^ CD3^+^ T cells/mL (depending on CD3^+^ T cell percentage of the respective sample).

b. Add 50µL HD-derived PBMCs per well (16 x 10^4^ effector cells per well).

c. Add 100µL 2ng/mL Blinatumomab per well (1ng/mL final concentration).

d. As a positive control for cell death: Add 100µL 20µM venetoclax to 100µL cell suspension (10µM final concentration).

e. Centrifuge the plate for 10 minutes at 200 x g.

f. Incubate the cells in a humified incubator (37⁰C, 5% CO_2_) for 24 hours.

5. Perform endpoint flow cytometry analysis.

a. After culture, resuspend the cells to dissociate the spheroids in order to ensure proper flow cytometry staining.

b. Transfer 100µL cell suspension per well to a non-sterile 96-well plate.

c. Add 10µL 0.1µM DioC6 solution per well.

d. Incubate the cells for 20 min at 37⁰C in the dark.

e. Optional: Perform flow cytometry surface staining.

i. Centrifuge cells for 5 minutes at 578 x g.

ii. Wash cells with 100µL 0.5% BSA diluted in PBS.

iii. Resuspend cells in 20µL antibody staining.

iv. Incubate cells for 30 minutes at 4⁰C on a plate shaker at 450 rpm in the dark.

v. Wash cells 2x with 100µL 0.5% BSA diluted in PBS

vi. Resuspend cells in 100µL 0.5% BSA diluted in PBS.

f. Add 5µL 0.2µM TO-PRO-3 solution per well.

g. Incubate the cells for 5 min at 4⁰C in the dark.

h. Measure cells on a flow cytometer.
